# Large-Scale Genomic Reorganizations of Topological Domains (TADs) at the *HoxD* Locus

**DOI:** 10.1101/116152

**Authors:** Pierre J. Fabre, Marion Leleu, Benjamin H. Mormann, Lucille Delisle, Daan Noordermeer, Leonardo Beccari, Denis Duboule

## Abstract

**Background:** The transcriptional activation of *Hoxd* genes during mammalian limb development involves dynamic interactions with the two Topologically Associating Domains (TADs) flanking the *HoxD* cluster. In particular, the activation of the most posterior *Hoxd* genes in developing digits is controlled by regulatory elements located in the centromeric TAD (C-DOM) through long-range contacts. To assess the structure-function relationships underlying such interactions, we measured compaction levels and TAD discreteness using a combination of chromosome conformation capture (4C-seq) and DNA FISH.

**Results:** We challenged the robustness of the TAD architecture by using a series of genomic deletions and inversions that impact the integrity of this chromatin domain and that remodel the long-range contacts. We report multi-partite associations between *Hoxd* genes and up to three enhancers and show that breaking the native chromatin topology leads to the remodelling of TAD structure.

**Conclusions:** Our results reveal that the re-composition of TADs architectures after severe genomic re-arrangements depends on a boundary-selection mechanism that uses CTCF-mediated gating of long-range contacts in combination with genomic distance and, to a certain extent, sequence specificity.

## BACKGROUND

Genes involved in key developmental processes are usually expressed in different tissues and at different times and hence they require particularly precise regulatory controls [1, 2]. To achieve this complexity in their transcription patterns, they often rely on the presence of multiple regulatory elements, including enhancer sequences (e.g. [3]. In addition, multiple enhancers can serve the same or a related specificity, either by acting as shadow enhancers to ensure robust transcription under adverse conditions [4-6], or by complementing one another in a ‘regulatory archipelago’ to integrate and assemble various parts of one large transcription domain [7].

In vertebrates, accumulating evidence suggest that most enhancers are located within regions of the genome initially referred to as a regulatory landscapes [8], often localized at some distance from the target gene(s). The genome-wide application of chromosome conformation capture (see e.g. [9]) has revealed the existence of Topologically Associating Domains (TADs), an intermediate level of chromatin domains wherein enhancer-promoter interactions seem to be restricted and privileged [10-12]. TADs tend to be evolutionary conserved [10, 13, 14] and are characterized by constitutive contacts, involvement of chromatin architectural proteins, and hence they can be observed both in transcriptionally active and inactive contexts. As a consequence, regulatory landscapes and their target gene(s) often overlap with TADs [15, 16]. However, the causality question as to whether TAD can restrict enhancer-promoter contacts or, conversely, how important enhancer-promoter contacts are in the building of a TAD, remains to be clearly established (see [2, 17]).

The interplay between multiple enhancers, constitutive contacts and both the nature and the extent of a particular TAD can be advantageously studied by using the *HoxD* gene cluster under physiological conditions *in embryo.* These genes are transcribed in distinct combinations during embryonic development in a tissue- and time-specific manner, following their regulation by a series of tissue specific enhancers [7, 16, 18]. As in many other contexts, the identification of these regulatory sequences relied either upon particular histone modifications, chromatin accessibility or on the use of chromosome conformation capture (4C). In these cases, several millions cells were used, thereby precluding the possibility to make a precise account of either the number of possible 3D structures or of the underlying dynamic processes.

During digit development, several long-range enhancers located in the centromeric TAD (hereafter referred to as C-DOM) are required to control the transcription of a set of target *Hoxd* genes, in particular of *Hoxd13* [19]. The targeted deletion of C-DOM almost entirely abrogated transcription in digits, whereas partial deletions gave intermediate outcomes suggesting that these so-called ‘regulatory islands’ are all required to achieve the final and full transcription specificity [7]. However, such sequences carry specific features and are not simply a sum of their parts, since the substitution of some regulatory islands by others through genomic rearrangements induced visible phenotypic consequences [20].

Here, we use the development of digits development to investigate whether various combinations of interactions occur in different cells. 4C studies can indeed lead to abusive interpretations as the averaging process may not reflect any of the individual cellular contexts (e.g. [21]). We also assess the importance of the distance *versus* sequence specificity of these regulatory islands towards target genes, by using a set of copy number variants including a series of nested deletions leading to important reorganizations of the C-DOM TAD. We report that the building of new TADs, after severe topological re-organizations, depends on both the presence of characterized specific or constitutive interactions, including potential CTCF-driven contacts, as well as on a relative distance effect, suggesting that intrinsic physical properties may also contribute to the shaping of these chromatin domains at this locus.

## RESULTS

### The C-DOM TAD as a functional compartment for digits enhancer sequences

The C-DOM involves the core interactions between *Hoxd* genes and their digits enhancers, as defined by the interaction profile of *Hoxd13,* the major target of these enhancers within the gene cluster. In order to assess the dynamics of these interactions, we initially evaluated in detail, the contacts established by *Hoxd13* over this TAD in both distal and proximal dissected limb bud cells. In distal autopod cells, *Hoxd13* is transcribed robustly whereas in proximal zeugopod cells, i.e. cells sharing a close developmental history, *Hoxd13* is inactive, thus allowing for a direct functional comparison (Fig. 1a). The examination of these 4C profiles (three different replicates) revealed the global map of *Hoxd13* contacts and allowed the identification of those interaction peaks that display the highest variation between cells where *Hoxd13* is active or inactive. In particular, contacts with islands III and IV were substantially increased in active distal cells, as well as a part of island II. On the other hand, most contacts between *Hoxd13* and the telomeric TAD (T-DOM) appeared more robust in inactive proximal cells (Fig. 1a).

**Figure 1:**
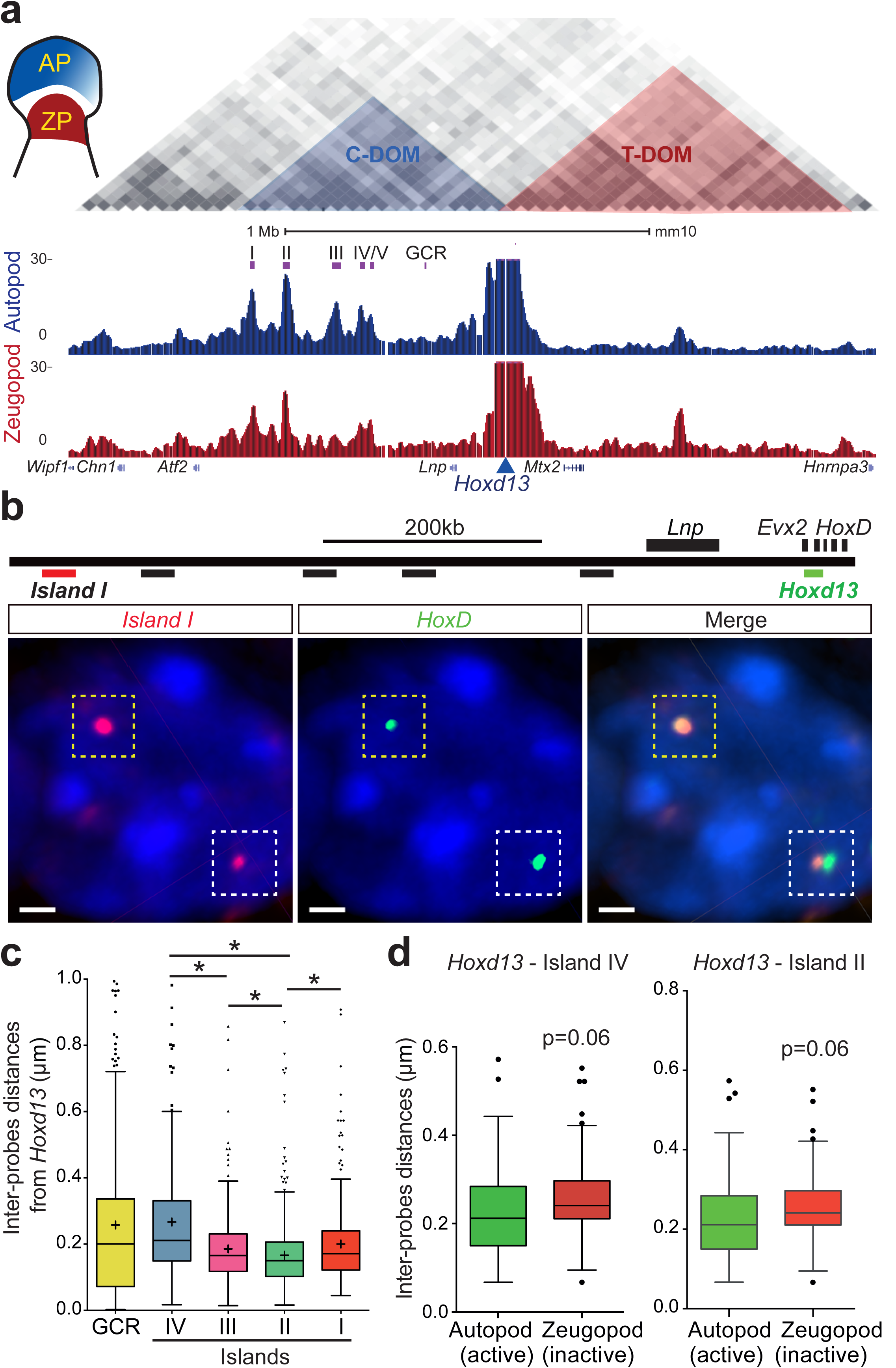
Interactions at the *HoxD* locus as seen by 3D DNA FISH and chromosome conformation capture (4C-seq). **a**, 4C interaction profiles (normalized signals) of *Hoxd13* in wild-type autopod (AP, blue) and zeugopod (ZP, red) cells, isolated from E12.5 embryonic mouse forelimb. The position of the TADs is highlighted on top (data from [10]). **b**, Position of fosmid probes used for 3D DNA-FISH. The images below show the FISH signals obtained for *Hoxd13* (green) and island I (red), as an example of both tightly associated signals (upper left allele, yellow dashed square) and separated signals (lower right allele, white dashed square). Scale bar: 1μm. **c**. Quantitation of inter-probe distances between *Hoxd13* and the regulatory elements in autopods ranked left to right, from the closest to the furthest in term of genomic distance. Kruskal-Wallis test was followed by Dunn’s multiple comparison test: * p<0.05. **d**, 3D DNA FISH distances as measured in autopod *(Hoxd13* active) and zeugopod *(Hoxd13* inactive) cells from E12.5 mouse forelimbs. Both Tukey boxplot representations show shorter distances in those tissues where *Hoxd13* is active.

In order to evaluate whether such dynamic variations in interactions were correlated with the position in the 3D space of each of the islands relative to *Hoxd13,* we performed 3D DNA-FISH in either distal or proximal cells (Fig. 1b) and found that most of these islands were located at a very short distance from *Hoxd13* (Fig. 1c).

While such close associations were not unexpected, due to the position of these islands within the same TAD, we nonetheless noticed that the interaction peaks identified previously [7, 16] and in Fig. 1a as displaying a dynamic behavior, were more closely associated with *Hoxd13* in positive, distal limb bud cells. In particular, islands II and IV were significantly closer to *Hoxd13* in active cells when compared to inactive proximal cells (Fig. 1d).

Also, the strongest associations as detected by 4C did not necessarily correspond to the closest genomic distances. Islands I and II for instance were often found within a 200nm distance from *Hoxd13* (66.5% and 74.5%, respectively), whereas the GCR sequence and island IV were scored further away (50.1% and 46.3% below 200nm) (Fig. 1b-c). Such differences in distances were significant (Mann Whitney test) and well supported by the strong interaction peaks detected on islands I and II in distal cells (Fig. 1a and [7, 16, 18, 22]). In addition, when we compared active and inactive cells, the spatial proximity between *Hoxd13* and its dynamic enhancers was decreased in the presence of active transcription, again supporting the results obtained using 4C (Fig. 1d **and Fig. S1**). Likewise, a shortening of the distance between the islands I and II themselves, specifically observed in distal cells, suggested that the regulatory landscape adopted a globally more condensed configuration, as shown previously for the entire C-DOM [23].

### Multipartite interactions

Of note, the distribution of distances observed displayed a great heterogeneity from cell to cell. Furthermore, the extent of variation was slightly different for each regulatory island suggesting that they may occasionally contact their target gene individually. Therefore, we performed DNA-FISH by combining three probes at the same time, either specific for three different islands, or for two islands as well as the *Hoxd13* target gene (Fig. 2a-b). We observed that several regulatory sequences can be detected juxtaposed in the same cell (Fig. 2a). These tripartite complexes however displayed heterogeneous configurations (Fig. 2a), despite the use of cells where the transcription of *Hoxd13* is supposed to be robust and homogenous [19]. Amongst the heterogeneous combinations scored, the occurrence of one island being located somehow at the interface between *Hoxd13* and another island was over-represented, suggesting that some of these genomic sequences may trigger the formation of larger regulatory structures.

**Figure 2:**
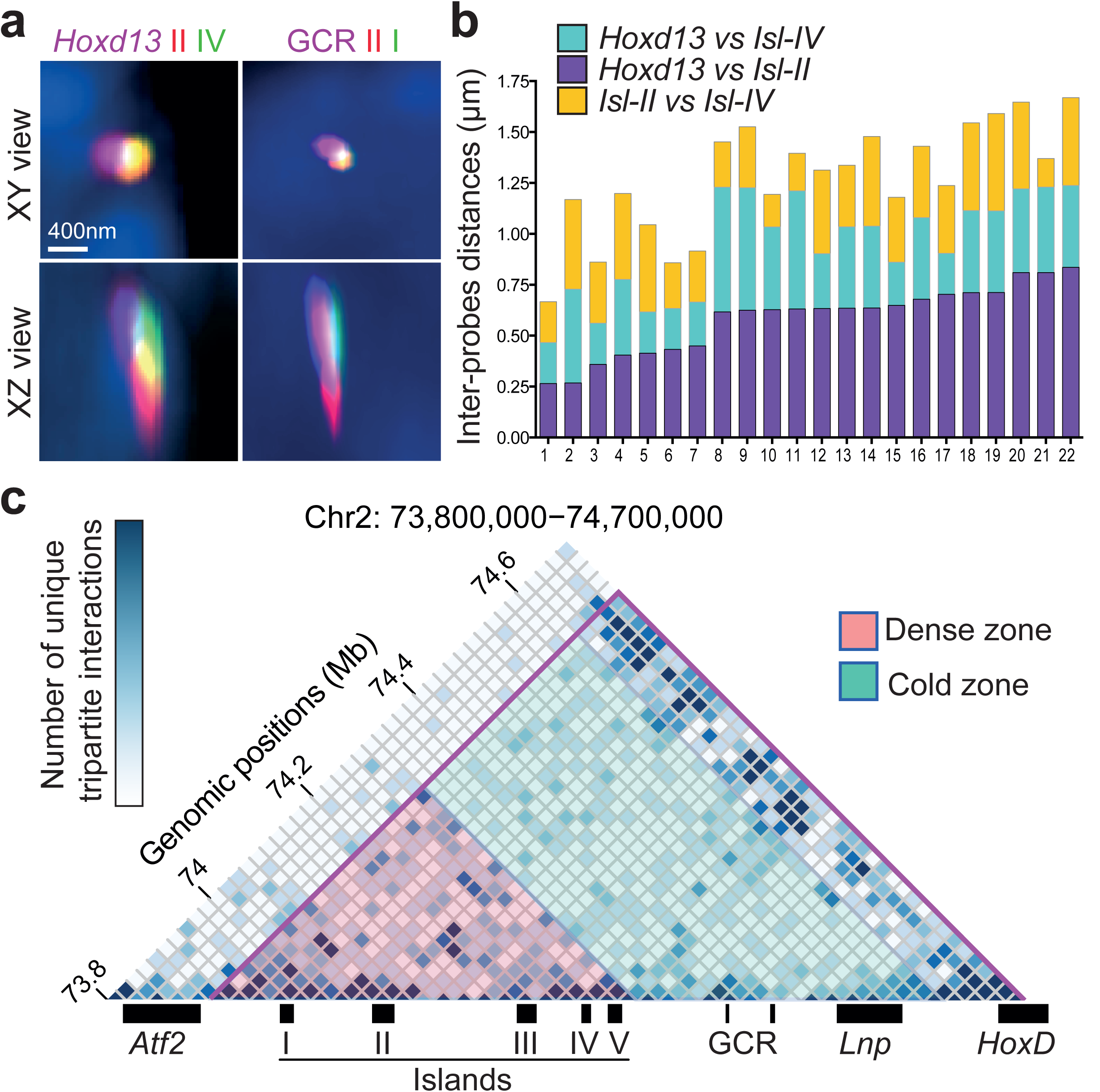
Tripartite interactions between *Hoxd13* and the regulatory islands. **a**, Orthogonal projections from 3D DNA-FISH (as in Fig. 1b) performed using either *Hoxd13* and two regulatory islands (left) or three islands (GCR, island II and island I). Scale bar: 400nm. **b**, 3D-DNA measurements showing variations in the distribution of physical distances between *Hoxd13* and island II (purple, bottom), *Hoxd13* and island IV (cyan, intermediate) or island II and island IV (yellow, top). **c**, Heatmap of unique tripartite interactions generated from 4C-seq data using *Hoxd13* as a viewpoint. The matrix is represented with a bin size of 20kb. The purple line shows the limits of the C-DOM as identified in [10].

4C-seq was recently used to identify multiple and concomitant interactions as well [24, 25]. We thus investigated whether the complexes observed by DNA-FISH could also be detected through tripartite interactions using 4C matrices (Fig. 2c). Such interactions between two regulatory islands, on the one hand, and *Hoxd13* on the other hand, were indeed scored. Their frequencies were significantly higher in a more distant ‘hot zone’ in the C-DOM TAD, where the five islands are concentrated. In this region, 67% of 20kb large bins were detected in at least one tripartite interaction (Fig. 2c; red zone), whereas a cold zone was observed closer to the *Hoxd13* target, where only 39% of bins were involved in tripartite contacts (Fig. 2c green zone). Altogether, these observations suggested that *Hoxd13* does not physically interact with all of its regulatory elements in every cell all the time. However, it can interact with multiple elements in the same cell at the same time.

### Regulatory *versus* genomic distances

From the results above, the genomic distance (as measured in base pairs) within C-DOM does not correlate with the ‘regulatory distance’ (as measured by the capacity to elicit a regulatory outcome). To determine to what extent the genomic distance can affect the regulatory contact, we assessed whether the relocation of a particular enhancer sequence at a distance outside C-DOM can abolish its contacts with the target gene. To this aim, we looked at a large inversion where the two more distal islands I and II were relocated 2.4 megabase (Mb) away (*HoxD^invTpSB1-Itga6)^* [20]. In this context, we analyzed the spatial distance between *Hoxd13* and island I by using 3D DNA-FISH and observed a clear loss of proximity as the distance between *Hoxd13* and island I was increased (Fig. 3b-c). This separation was similar to distances previously observed between other segments of DNA that do not form functional gene-enhancer contacts [26].

The 4C interaction profiles obtained using this large inversion showed a near complete absence of contact between either island I or island II with *Hoxd13,* confirming this increased distance (Fig. 3d-3f; RegY). This suggests that, to some extent, a minimal genomic distance may be required to allow and stabilize an interaction. However, increased contacts were observed in other instances, over much larger distances for instance when using the 28Mb large *HOXD^Inv(HoxDRVIII-Cd44)^* inversion [27]. In this case indeed, a faint interacting region localized 19Mb far from the *HoxD* cluster (the *Alx4* gene promoter; [26]) was re-positioned at a distance of 9.2Mb from *HoxD,* leading to a visible decrease in distance by 3D DNA-FISH (**Fig. S2a-b**) associated with a robust increase in the corresponding 4C signals (**Fig. S2c**).

In the case of the *HoxD^inv(TpSB1-Itga6)^* inverted allele, however, the loss of 4C contacts between island I and *Hoxd13* was perhaps compensated by novel DNA contacts between *Hoxd13* and the sequences relocated at the position of the displaced islands (Fig. 3d-e; RegZ). The novel and ectopic contacts were discrete and indeed of intensities comparable to those of known interactions normally occurring within this TAD. This result suggested that C-DOM had been re-organized as an interaction domain of a similar size. This internal TAD re-organization, however, substantially impacted the contact dynamics of those islands that had not changed their genomic distances to the target gene, in particular islands III and IV, which displayed reduced peak sizes in the inverted allele (Fig. 3d; red star), whereas island V showed the opposite effect giving rise to a global profile that resembled more that obtained with proximal rather than distal, limb bud cells (Fig. 1a and 3d).

However, the extent of the contacts gained by *Hoxd13* with naive DNA sequences coming from the inversion did not depend upon the genomic distance but instead, involved some sequence specificity. Indeed when we compared these additional contacts (Fig. 3; RegZ) with those gained after using the *HoxD^Inv(Nsi-Itga6)^* allele, which inverted and thus re-localized the entire C-DOM [16] (Fig. 3e), we observed that in both cases the gained interactions span about 200kb of the inverted DNA, with particularly strong peaks mapping on the promoter of *Rapgef4,* a gene that was brought to the vicinity of the *HoxD* locus by both inversions. Therefore, in both genomic rearrangements, this gene acted as a landmark in the building of new interactions domains, regardless of the overall sizes of these new domains.

**Figure 3:**
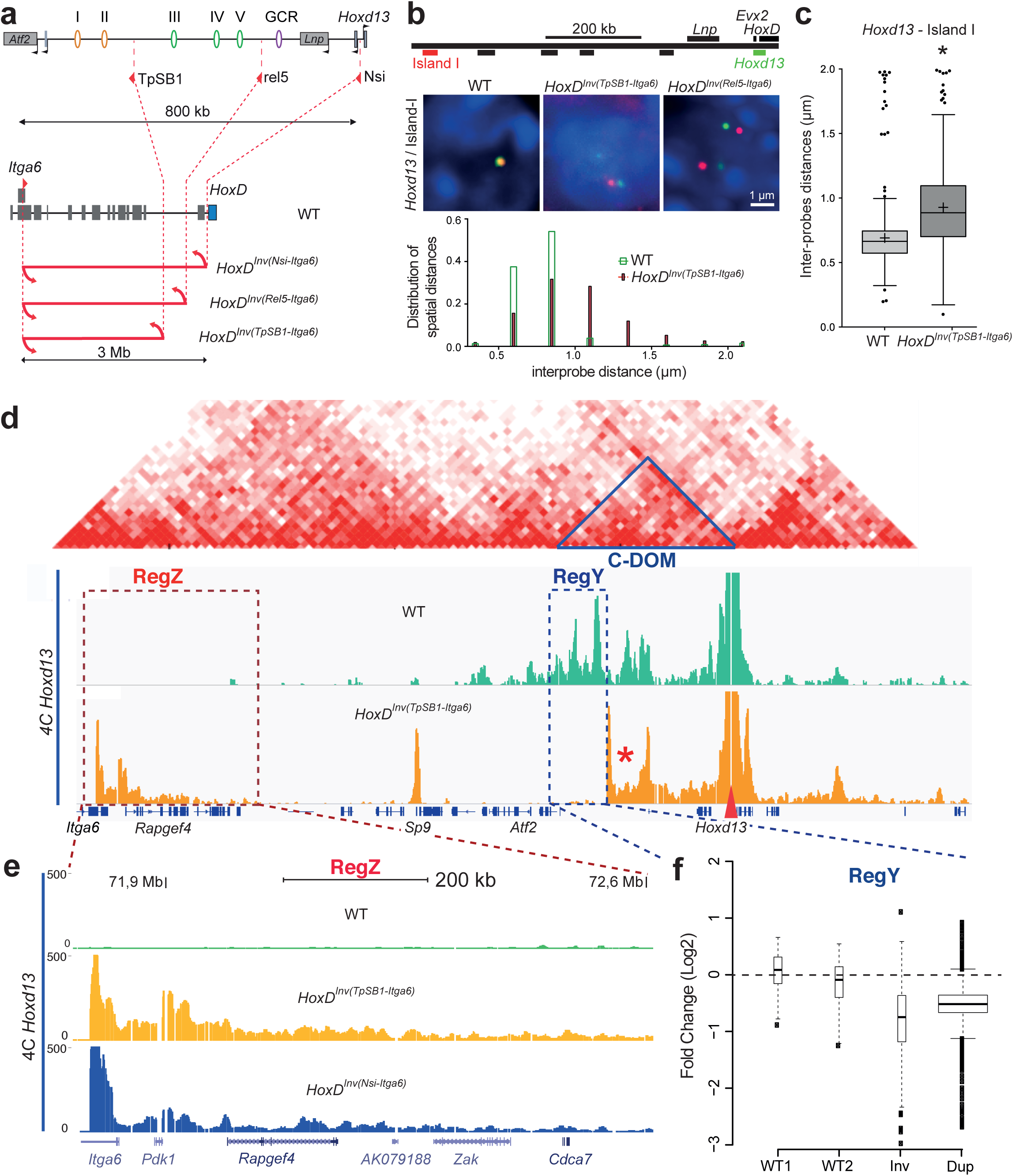
Impact of increasing distances upon interactions with *Hoxd13.* **a**, Schematic of the large inversions displacing regulatory islands away from *Hoxd13.* **b,** DNA-FISH signals for *Hoxd13* (green) and island I (red) in distal limb bud autopod cells dissected from embryos mutants for either one of two large inversions. Scale: 1 μm. Below are shown the distribution of spatial distances in the WT (green) and the inverted allele (purple). **c**, Quantification of 3D distance showing the increase in distance observed in **b** (Mann Whitney test, p<0.001). **d**, 4C-seq profile using *Hoxd13* as a viewpoint either in control (green), or the 2.4Mb large *HoxD^Inv(TpSB1-Itga6)^* inversion (orange), are aligned below the heatmaps generated using data from [10] with C-DOM indicated by a blue triangle. The *Hoxd13* viewpoint is indicated with a red arrowhead. **e**, Close-up of region Z where contacts decrease, in control and both *HoxD^Inv(TpSB1-Itga6)^* and *HoxD^Inv(Nsi-Itga6)^* mutant samples. **f**, Quantitation of contacts mapping into region Y, with the same DNA sequence located at various distances from the target *Hoxd13* gene. Quantitations are shown either for two different control replicates, the 2.4Mb inversion (inv for *HoxD^In^*^v*(TpSB1*-*Itga6)*^) and a previously reported duplication *(HoxD^Dup(Nsi-TpSB1)^* or ‘dup’) where this particular region is moved 300kb further away from *Hoxd13* [20]. A higher number of fragments included in the analysis for the duplication (due to a slightly different 4C method, on chip) is shown by the thicker boxplot.

### Impact of TAD rearrangement on transcription

To assess whether these recomposed TADs were still able to regulate their *Hoxd* target genes, we measured the expression of posterior *Hoxd* genes using RT-qPCR (Fig. 4a-b). All *Hoxd* genes previously shown to interact with C-DOM [7] displayed somewhat reduced steady-state levels of mRNAs (Fig. 4b). We next used RNA-seq to more globally evaluate the transcription of genes located in *cis* of the *HoxD* cluster, on a large scale and could confirm the decreased transcription of posterior *Hoxd* genes in these re-organized topologies (Fig. 4c). In addition, we detected a set of up-regulated genes (Supplementary table 1), including *Dlx1* and m*Dlx2*, two genes located a few Mb far from the *HoxD* cluster and neighboring islands I and II after their topological relocation following the inversion (Fig. 4d-e). We confirmed the apparent transcriptional up-regulation of *Dlx1* by RNA-FISH and observed a clear increase in signal intensity specifically in distal forelimb cells. In cells from the retina, a tissue where *Dlx1* is already expressed [28] but where the regulatory islands I and II are normally not functional, this transcriptional boost was not scored (Fig. 4f).

**Figure 4:**
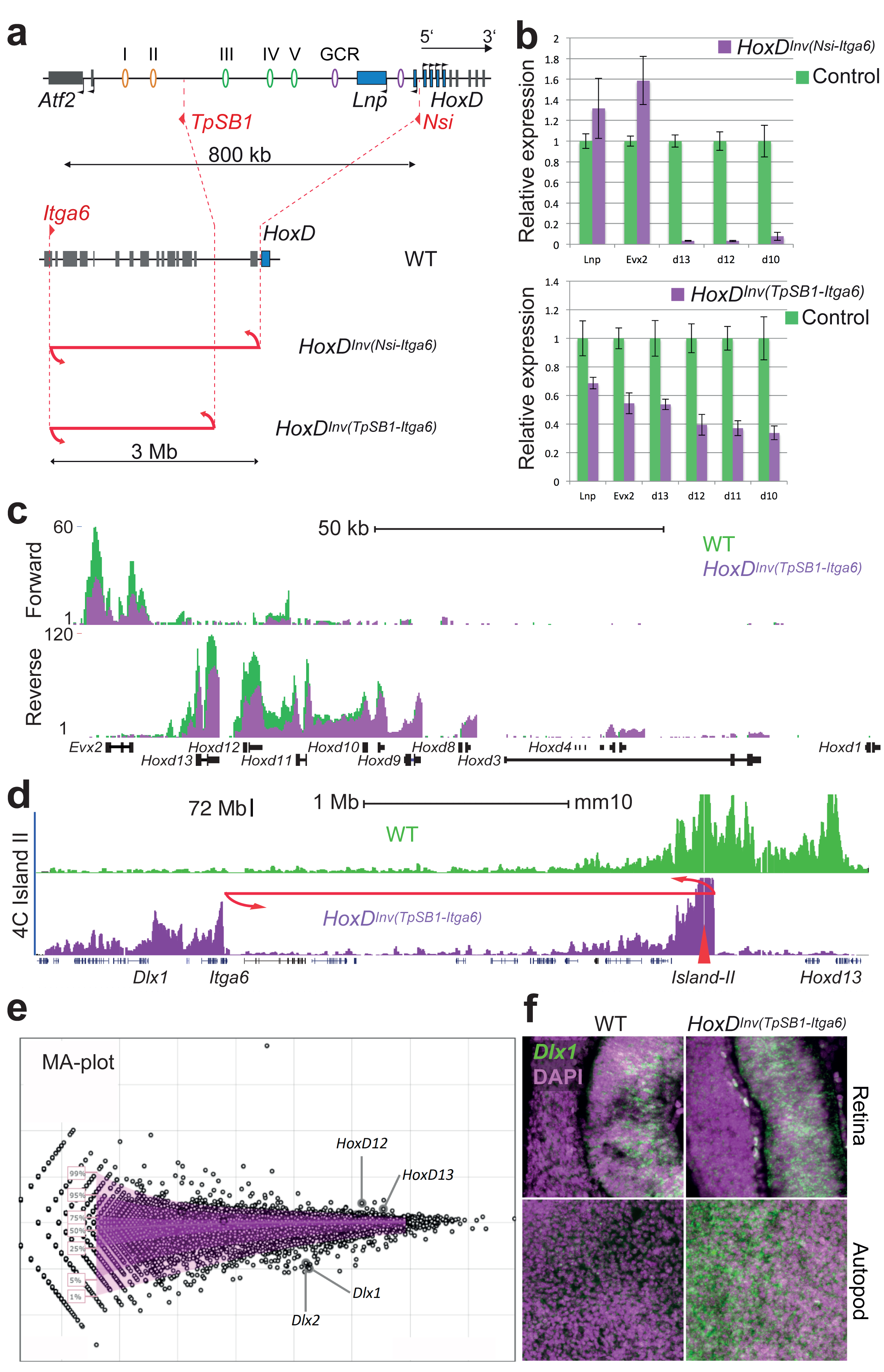
Impact of large inversions on transcription. **a**, Schematic of two large inversions displacing regulatory islands away from *Hoxd13.***b,** RT-qPCR quantitations of target *Hoxd* gene mRNAs in presumptive digit cells either from the inversions (purple) or from control cells (green) show a significant loss of *Hoxd10* to *Hoxd13* transcripts in both inversions. Error bars represent SD (n=3), * p<0.05, ** p< 0.01. **c**, Strand-specific RNA-seq profiles showing reduced transcription when the 2.4Mb large *HoxD^Inv(TpSB1-Itga6)^* inversion was used (green, WT; purple, inverted allele). **d**, 4C-seq profiles using Island II as a viewpoint (red triangle) in autopod cells from WT (green) and *HoxD^Inv(TpSB1-Itga6)^* inversion (purple) showing the gain of contacts with the *Dlx1* locus in the inversion allele. **e**, Changes in expression as measured by RNA-seq are represented as a MA Plot, with *Hoxd12* and *Hoxd13* down-regulated in the inversion (as in **b**) whereas *Dlx1* and *Dlx2* were up-regulated. *Dlx1* up-regulation is controlled by RNA-FISH in **f,** with the upper panels showing the retina and the lower panel distal limb autopod cells (both in E12.5 embryos).

### Pre-determined reallocation of contacts

The reallocation of contacts observed between *Hoxd13* and non-related DNA after the large inversion suggested that genomic distance represents a physical constraint in the building of the C-DOM TAD structure. Alternatively, the inversion may lead to the redistribution of TAD ‘boundary’ elements, which would then redesign the TAD landscape. To evaluate whether removing specific parts of the C-DOM TAD resulted either in an increase of contacts over the remaining regulatory islands, or in the extension of the TAD to include novel points of interaction, we used a series of internal deletions (Fig. 5; *del-1* to *del-4).* We compared the 4C profiles of the various deleted configurations (Fig. 5 **and S3**) and noted a high conservation of sequence specific interactions, regardless of which elements of the TAD had been removed (Fig. 5b). Contacts persisted on the remaining islands independently from the change in genomic distances, indicating that the TAD structure in itself does not seem to be required for all internal contacts to be optimally established.

**Figure 5:**
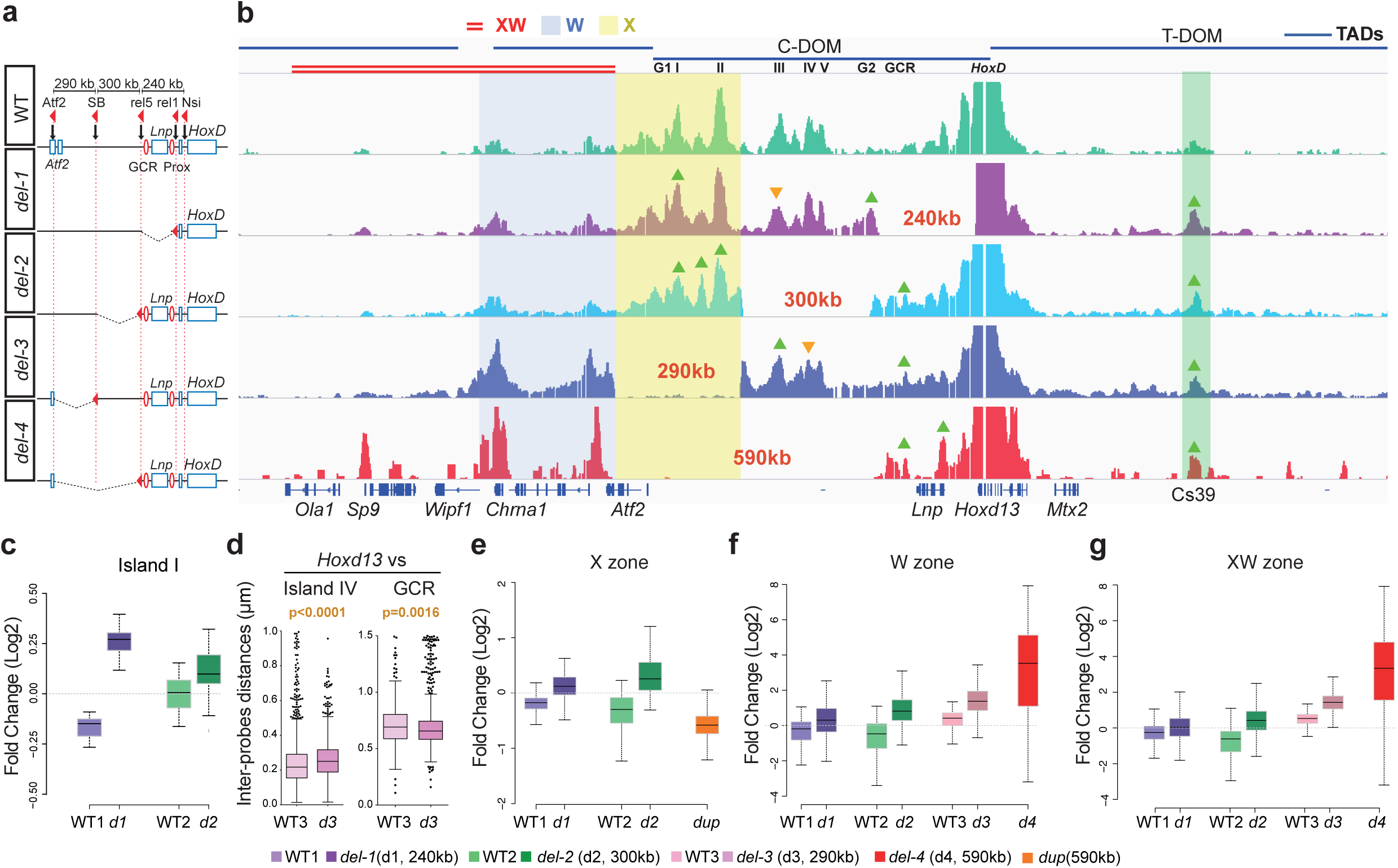
Reallocation of long-range contacts in reorganized TADs. **a**, Schematic of the deletion series showing the loxP site position used to generate the deletion (red arrows). **b**, 4C interaction profiles from *Hoxd13* as a bait in control (wild-type, green), *del-1* (purple), *del-2* (light blue), *del-3* (dark blue) and *del-4* (red). The four horizontal bars on top represent the TADs (as characterized by [10]. The position of the islands is indicated below the bar of C-DOM. The X zone is highlighted in yellow, the W zone is highlighted in blue and the double red line on top indicates the XW zone. The green triangles indicate statistically significant increases of contacts and orange triangle decreased contacts. **c,e-g**, The Y-axis represents a log2 scale. **c**, Fold change on specific contacts as observed in **b** for island I. **d**, Distance measurements from 3D DNA-FISH between *Hoxd13* and island IV (left) and GCR (right) in autopods from wild-type and *del-3*. **e**, Fold change in the region X (highlighted in yellow in panel **b**). **f**, Fold change in a region termed “W zone”, highlighted in blue and centromeric to all deletions. **g**, Fold change in a region termed “XW zone”, a 590kb large region highlighted in red, containing the W zone and extending further centromeric to all the four deletions.

In the *del-1* allele, shortening the distance between the islands and their target genes by 240kb had little effect on their interaction profiles (Fig. 5b). Statistical analyses however revealed that when islands I and II were positioned at the places of islands III and IV through the *del-1* deletion, their interaction peaks increased in intensity (Fig. 5c **and S3b**). Instead, the island III peak, which was shown to be specific to digit cells [16] was reduced even though located closer to its interacting genes in term of genomic distance (**Fig. S3c**). Such slight modifications in peak intensities were also observed in the *del-2* and *del-3* deletions (Fig. 5b; green and orange arrowheads and Fig. 5c-e; log2 fold change compared to their wild type littermates).

Interestingly, whenever a particular deletion induced the strong re-enforcement or even the appearance of an interaction peak not detected in control cells, this sequence usually corresponded to a region of specific interaction in another cellular context or, alternatively, in another TAD. The first case was best illustrated by the *del-1* allele, where interactions involving *Hoxd13* were increased with the GT1 and GT2 sequences (**Fig. S3c-d**). These two enhancers normally interact with this gene but only in the developing genitals and not in developing digit cells [18]. They were however recruited in the novel regulatory topology induced by this particular deletion.

The second case was observed with the *del-3* and *del-4* mutant alleles. The *del-3* deletion triggered *Hoxd13* to establish two major contacts upstream of its usual proximal TAD boundary, within what is the adjacent TAD in the wild type situation (Fig. 5b; region W). These two interactions peaks however could already be observed in the control and *del-1* and *del-2* profiles, where the TAD boundary is not deleted, though with a much lower intensity. Therefore, deletion of the TAD border merely reenforced the contacts, which were already established yet at a much lower frequency (Fig. 5f). This was supported by the largest internal *del-4* deletion, which removed regulatory islands I to V. In this case, the severely decreased expression of *Hoxd13* reported for this configuration [7] correlated well with the robust interaction with the *Sp9* gene. Indeed these loci were shown to interact with one another when transcriptionally inactive [26]. There again, weak interactions with this gene were already scored in control cells, suggesting that these interactions were not fully *de novo* induced by either reducing the distance or deleting a TAD boundary. However, the contacts extended further in the surrounding region XW until the gene *Ola1* (Fig. 5b).

In the *del-3* allele, we noted that the strong increase in contacts in region W occur even though the genomic distance is the same as in the *del-2* allele, where such gains of contacts were not observed. This was likely due to the removal of the proximal TAD boundary in the *del-3* allele, whereas this boundary is still present in the *del-2* chromosome. In this view, the TAD boundary seems to be important to properly assign interaction strengths to contacts that normally occur, sometimes at very low frequencies. The strong distal interactions in *del3*, leading to what appears to be a fused TAD including C-DOM and region W (Fig. 5b), also slightly re-organized contacts occurring in the proximal part of the TAD such as an increase in contacts with island III and a decrease with island IV. There again, however, this reorganization involved sequences already interacting, rather than de novo contacts (Fig. 5b).

Finally, any genomic modification of the C-DOM landscape led to an increase of contacts between *Hoxd13* and the TAD located at the opposite side of the gene cluster (T-DOM). This was particularly visible with one sequence referred to as Cs39, an enhancer sequence active in proximal cells, during the development of the zeugopods [16, 29] (Fig. 5b; green bar). There again, this region normally already contacts *Hoxd13* at low levels (Fig. 1a and Fig. 5b, upper 4C profile). This observation suggested that the optimal topological configuration to transcribe *Hoxd13* in digit cells involves the presence of a native C-DOM, as all deletions tested lead to decreases in mRNA levels. In this optimal situation, *Hoxd13* is fully engaged into interacting with the centromeric islands. Whenever a genomic perturbation is induced, *Hoxd13* looses some interactions with C-DOM and thus can be re-directed towards the major T-DOM interactions, concomitantly to a loss of transcriptional outcome.

### Boundary effects

In the *del-4* allele, where all five islands were deleted, not only did the contacts extend into the neighboring TAD (Fig. 5b,f; zone W), but further continued up to a distance similar to the genomic segment that had been deleted (Fig. 5b,g; zone XW), thereby reaching the *Sp9* gene and behind. In the *del-1* and *del-2* alleles, however, the extension of the interactions towards the centromeric side importantly decreased at the *Atf2* gene and, in the *del-3* allele, contacts increased up to the *Chrna1* gene yet not really behind (Fig. 5b). Regarding the *del-1, del-2* and *del-3* cases, these observations could be attributed to the presence of a potential TAD boundary, as previously mapped by Hi-C [10]. Indeed the *del-1* and *del-2* alleles did not physically affect such a boundary, whereas the *del-3* allele removed the TAD boundary between C-DOM and TAD V, allowing for *Hoxd13* contacts to extend into the latter TAD.

However, the results obtained with the *del-4* allele could not be interpreted in this way since no additional boundary was deleted when compared to *del-3,* yet the interaction profile largely extended in the centromeric region, reaching the *Sp9* and the *Ola1* gene body (Fig. 5b). Also, in the two large inversions discussed above (Fig. 3e and 4a) where the C-DOM TAD boundary was removed in both cases, the contacts extended up to the same gene *Rapgef4.* Because *Hoxd13* is located close to a TAD boundary and is surrounded by CTCF sites [30], which are associated with TAD boundaries [10, 31, 32], we looked at the presence of bound CTCF in these various regions to try and explain these differences.

We performed ChIP-seq for CTCF using the autopod from E12.5 embryo and confirmed the binding pattern previously reported by a ChIP-on-Chip approach using the same embryonic tissue [30]. We also compiled the ENCODE data for CTCF and the cohesin subunit RAD21 and all datasets revealed five CTCF peaks surrounding the *Chrna1* locus, with four of them being also bound by cohesin (Fig. 6a-b), further defining this region as a potent TAD boundary and explaining the re-organized interaction profiles observed with the *del-3* alleles, as well as the weak contacts already scored between *Hoxd13* and this region in the *del-1* and *del-2* alleles.

**Figure 6:**
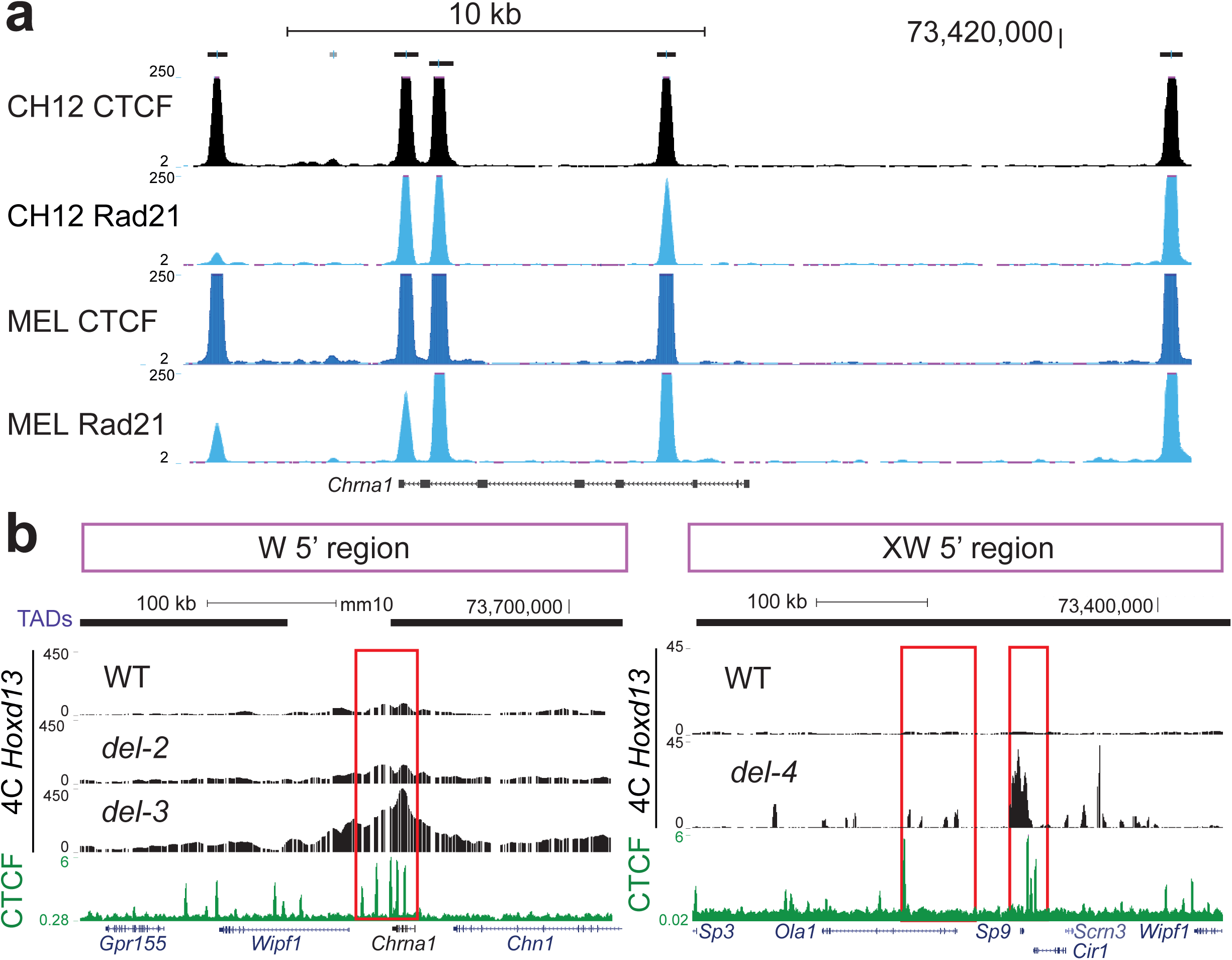
Re-organized TAD boundaries are CTCF-rich regions. The tracks corresponding to the CTCF and cohesion Chip-seq data are either extracted from ENCODE (**a**), or from our distal limb bud cells at E12.5 (**b**). They are displayed in the UCSC Genome Browser (http://genome.ucsc.edu). The regions emphasized are the DNA segments where contacts drastically decrease, as observed in Fig. 5b.

Multiple sites of bound CTCF were also scored at the *Sp9* locus (3 peaks) and near *Ola1* (1 peak), despite the fact that this zone was never reported to be a TAD boundary (Fig. 6b). Finally, we observed a comparable enrichment of CTCF peaks (≥7) in both the promoter and the gene body *Rapgef4* (not shown). Therefore, while it seems that under our physiological conditions, the reorganization of topological domains after targeted CNVs will generally involve the use of bound CTCF as novel landmarks to exert a boundary effect, exceptions exist such as the *del-3* allele, where the *Chrna1* locus, a strong boundary element is ignored and interactions established between *Hoxd13* and *Sp9,* which itself lies in the middle of another TAD. In this particular case, the genomic distances as well as potential internal constraints associated should be carefully considered.

## DISCUSSION

### Multiple interactions

Our 4C-seq approach identified multiple contacts between more than two sequences at a time, suggesting that the overall interaction profile observed by using millions of cells at least partially represents what is happening within single cells. It nevertheless does not allow us to propose any conclusion regarding the overall dynamics of contacts within this particular C-DOM, i.e. whether the increased occurrence of particular multiple contacts is due to a longer interaction time or, alternatively, to a higher frequency of contacts between the sequences involved. Therefore, while these results suggest that some contacts may be cooperative or perhaps requested to properly bring an enhancer at the vicinity of its target gene, the number of potential TAD configurations and their respective frequencies (see [21]) cannot be predicted. Such tripartite interactions at this locus were also studied recently in ES cells where transcription is not observed and may thus reflect a constitutive state. In this latter set-up, however, the tripartite interactions were more homogenous and the cold zone was less defined than in digit cells [33].

This cold zone we identified in our tripartite 4C matrices corresponds to a region of low contacts when using the *Hoxd13* locus as a viewpoint. This was confirmed by our FISH data, where a sequence falling within this space, the GCR, showed the highest mean distance to *Hoxd13* (Fig. 1c). Also, a previous serial deletion analysis *in vivo* had suggested that each part of this TAD was functionally rather independent from one another [7]. Altogether, by using several parameters, the TAD sub-region covering islands I to V appeared more structured than the region immediately flanking the *HoxD* cluster in distal limbs.

### Contact re-allocations

Noteworthy, out of our numerous analyzed mutant alleles that re-compose TAD structure, none of them produced a strong interaction that was not previously observed either in a control, or in another mutant situation, nor did they radically suppress any existing interaction. For example, when both large inversions were used, none of the contacts left between *Hoxd13* and the remaining parts of the TAD disappeared, despite the re-composition of the structure due to the addition of foreign DNA (Fig. 3). Likewise, when various deletions including parts of the TAD were analyzed, none of the remaining interactions were seriously affected, even when large deletions were used. The *del-2* allele for instance removed three strong points of contacts with *Hoxd13* (islands III, IV and V), yet the interaction profile of *Hoxd13* with the remaining islands I and II was almost as in control limbs. We take this as a strong indication that contacts within a TAD have only a moderate impact upon the global architecture of this TAD.

An exception to this was the behavior of island III, a *Hoxd13*-interacting sequence exclusively detected in developing digits (e.g. [29]). In the *del-1* allele where island III became closer to the cluster due to the deleted piece of DNA, its interaction with *Hoxd13* was importantly diminished (Fig. 5b **and Fig. S3c**). This effect was also observed in the large inversion whereby islands I and II had been displaced (Fig. 3d). In these genomic contexts, the loss of island III contacts paralleled the loss of activity of *Hoxd13,* suggesting that part of the functional impact of these re-organizations was due to *in cis* effects on non-deleted sequences, rather than solely to the deleted sequences themselves.

Of particular interest also was the appearance, in the *del-1* context, of the two GT1 and GT2 interaction peaks, not observed in control digit cells but present in the developing genitals where *Hoxd13* is expressed equally strongly (Fig. 5b and **Fig. S3**)[18, 34]. Therefore, the re-organization of the topological domain resulted in the recruitment of sequences that are used in a different context to control the same target gene. These results suggest that potential sites of interactions are fixed, but that tissue-specific TAD architectures can select a defined sub-set for regulatory use. Unfortunately, we could not directly assess whether these two sequences actively participated to the transcriptional outcome of the domain or alternatively, whether these contacts were purely induced by the new topology adopted after the *del-1* deletion without any particular effect.

### TAD modularity

The various regulatory islands do not seem to require a domain structure to exert their functional potential. They need not be embedded into a TAD to properly work, as shown both by deletion and transgenic analyses [7, 18], suggesting that TADs at *Hox* loci have a functionally modular structure. This was confirmed by functional analyses of some of the mutant configurations used in this work. For example, the transposition of islands I and II close to the *Dlx1/Dlx2* locus outside of their ‘native’ TAD, lead to the up-regulation of the latter two genes in a domain where they are normally not transcribed. Such regulatory side-effects [8] can often be observed after large *in-cis* (see [35] or *in-trans* [36] genomic re-arrangements. As the *Dlx1-Dlx2* locus is normally covered by polycomb complexes in this cell type [29], the contacts with enhancer sequences may help evicting these repressive complexes, as described in the globin system [37]. Aberrant transcriptional outcomes deriving from such CNV-dependent enhancer-promoter re-allocations are likely to cause genetic diseases in a variety of instances [38-40].

### TAD boundaries, CTCF and pre-set interactions

The analyses of our mutant genomes where the centromeric *HoxD* TAD is reconfigured in different ways revealed two trends, which may reflect more general properties of chromatin folding. The first is that whenever strong interactions were gained between *Hoxd13* and sequences located outside the C-DOM, as a result of a deletion, these sequences already displayed some (very) weak interactions in the control situation. Therefore, these ectopic contacts were not *de novo* interactions but rather the re-enforcement of pre-existing weak interactions, established despite the presence of the original TAD boundary. The deletion of such a TAD boundary, for example in the *del-3* allele, induced a strong leakage of *Hoxd13* interactions but specifically towards sequences already weakly contacted in control cells. We take this as an indication that the contact map of a particular gene is likely independent from its TAD environment. In this view, TADs impose a bias in high-affinity contact distribution by favoring local interactions within a spatial framework over outside contacts.

The re-organization of TADs often involves the presence of architectural proteins such as CTCF and cohesin [38, 41, 42]. Here again, either in the *del3* allele or in our large inversions, the ectopic contacts were gained up to a region rich in such proteins, as in the case of *del-3,* where *Hoxd13* interactions could extend up to the *Chrna1* locus, which clearly acted as a new TAD boundary. However, in the *del-4* configuration (a deletion larger than *del-3* but containing the same boundary elements), contacts extended beyond the *Chrna1* locus even though it was bound by CTCF at five positions, four of which also contain cohesin, to reach *Sp9,* another locus with bound CTCF. In this context, increased contacts with Sp9 may also be facilitated by the H3K27 trimethylation of the inactive *Hoxd13* and thus reflects the polycomb-associated contacts [26]. We consider this as an indication that boundary elements are not sufficient to impose a TAD structure and that other parameters may be equally important in shaping chromatin at this structural level. This latter result, along with our observations on the two large inversions, suggest that the physical distance may be important, as TAD re-organization at least at this locus tend to generate interactions profiles of rather comparable sizes, perhaps reflecting intrinsic forces or constraints at work at this scale of the chromatin fiber. This may support the idea, based on FISH data, that the chromatin fiber has a random-walk configuration but confined within a defined volume in the range of the megabase [43-45].

## METHODS

### Animals

Mice were handled according to the Swiss law on animal protection (LPA), with the requested authorization (GE/81/14 to D.D.). Mice were raised and sacrificed according to good laboratory practice standards. Tissues were isolated from E12.5 embryos, either wild-type or mutant for four different deletions as well as two large inversions. The deletions were *HoxD^Del(rel1-rel5)^* (referred to as *‘del1’* throughout the paper), *HoxD^Del(rel5-TpSB1)^* or *del2*, *HoxD^Del(TpSB1-Atf2)^* or *del3* and *HoxD^Del(rel5-Atf2)^* or *del4* [7]. The inversions were *HoxD^Inv(TpSB1-Itga6)^* and *HoxD^Inv(Nsi1-Itga6)^* [20].

### Statistical analysis

For DNA FISH analyses, the differences between samples were evaluated with the Kruskall-Wallis test, followed by Dunn’s multiple comparison post-test. For the 4C-seq boxplots (Fig 1, Fig 5 and S4), the statistical differences between islands and regions of interest (islands, GCR, Cs39, *Chrna1* and the three zone described) were assessed by using pairwise Wilcoxon Rank Sum Tests, followed by BH corrections for multiple testing (FDR, [46]).

### 3D DNA-FISH

3D DNA fluorescent *in situ* hybridization was conducted as previously described [26, 47]. E12.5 mouse embryos were fixed in 4% paraformaldehyde, embedded in paraffin blocks and cut at 6 μm. Sections were oriented such that cells belonging either to the distal (autopod) or proximal (zeugopod) parts of the growing limb bud could be unambiguously discriminated. Probes were prepared by nick-translation with either directly labeled nucleotides (Ulysis alexa 647, Life Technologies) biotin- or digoxigenin-UTP using fosmid clones obtained from the BACPAC Resources Center (https://bacpac.chori.org/) and listed in Supplementary table 2. 100 ng of DNA were used with 7 μg of Cot1-DNA and 10 μg of sonicated salmon sperm DNA. They were labeled using either digoxigenin- or biotin-dUTP by nick translation with fluorescent revelations as described previously [47], using either Alexa 647, Alexa 568 or Alexa 488 as fluorophores. Slides were stained with DAPI and mounted in ProLong Gold (Life Technologies). Images were acquired using a B/W CCD ORCA ER B7W Hamamatsu camera associated with an inverted Olympus IX81 microscope. The image stacks with a 200 nm step were saved as TIFF stacks. Image reconstruction and deconvolution were performed using FIJI (NIH, ImageJ v1.47q) and Huygens Remote Manager (Scientific Volume Imaging, version 3.0.3). Distance measurements between probes signals were determined using an automated spot/surface detection algorithm followed by visual verification and manual correction using IMARIS version 6.5, Bitplane AG and Matlab 7.5, MathWorks SA. Data from figure 1 were evaluated only using manual measurements. Statistical significance analyses of distances were performed using Mann-Whitney test (in Fig. 1d; 3c; 5d; **S1b** and **S2b**), or using the Kruskal-Wallis test followed by Dunn’s post test (in Fig. 1c).

### RNA-FISH

E12.5 distal limbs were fixed in 4% paraformaldehyde, 15% sucrose and then frozen in OCT. 25 μm cryostat sections were dried for 30 minutes, post-fixed in 4% paraformaldehyde for 10 minutes and quenched with 0.6% H2O2 in methanol for 20 minutes. Slides were then processed using the Ventana Discovery xT with the RiboMap kit. The pretreatment was performed with mild heating in CC2 for 12 minutes, followed by protease3 (Ventana, Roche) for 20minutes at room temperature. Finally, the sections were hybridized using automated system (Ventana) with a *Dlx1* probe diluted 1:1000 in ribohyde at 64°C for 6 hours. Three washes of 8 minutes in 2X SSC followed at hybridization temperature (64°C). Slides were incubated with anti-DIG POD (Roche Diagnostics) for 1 hour at 37°C in BSA 1% followed by a 10 minutes revelation with TSA substrate (Perkin Elmer) and 10 minutes DAPI. Slides were mounted in ProLong fluorogold. Images were acquired using the same procedure as for DNA-FISH.

### RNA-seq

E12.5 distal forelimbs were dissected and isolated using Trizol LS reagent (Life Technologies) to generate total RNA tissue samples. RNA-Seq was performed according to the TruSeq Stranded Illumina protocol, with polyA selection. The strand-specific total RNA-seq libraries were constructed according to the manufacturers instructions (Illumina). Sequencing was done using 100 bp single-end reads on the Illumina HiSeq system according to the manufacturer’s specifications. RNA-seq reads were mapped to ENSEMBL Mouse assembly NCBIM37 and translated into reads per gene (RPKM) using the RNA-Seq pipeline of the Bioinformatics and Biostatistics Core Facility (BBCF) HTS station (http://htsstation.epfl.ch). RNA-seq data can be found in the Gene Expression Omnibus (GEO) repository under accession number GSE_______.

### 4C-seq

Micro-dissected E12.5 proximal or distal limb bud tissues were dissociated, fixed with 2% formaldehyde, lysed and stored at -80°C. The nuclei from 10 pairs of distal or proximal forelimbs were then digested with a sequence of *NlaIII* and *DpnII,* followed by amplification according to [48]. The ligation steps were performed using high concentrated T4 DNA ligase (M1794, Promega) and the inverse PCRs for amplification were carried out using primers specific for the various viewpoints [49]. *Hoxd13* amplification primers were previously described [49].

PCR products were multiplexed and sequenced with a 100bp single-end Illumina HiSeq flow cell. Demultiplexing, mapping to the mouse assembly GRCm38 (mm10) and 4C-Seq analysis were performed using the BBCF HTSstation (http://htsstation.epfl.ch and [50]), according to previously described procedure [49]. Briefly, the 4C-Seq fragments directly surrounding the viewpoints (2kb) were excluded for the rest of the analysis. Fragment scores were normalized to the total number of reads mapped and smoothed (running mean with a window size of 11 fragments). For comparison purpose, the 4C-seq profiles were normalized to the mean score of fragments falling into a region defined as the bait coordinates +/-1Mb. For quantification of 4C-seq profiles in specific islands or regions of interest (boxplots in **Fig. S1** and Fig. 5), the smoothed data, with or without profile correction were used. When appropriate (e.g., signals in Fig 1b), replicates were combined by averaging the resulting signal densities. In Fig 5 **and S3**, quantitative log2 ratios were calculated by dividing the fragment scores with the means in WT1, WT2 and WT3. 4C-Seq data is available from the Gene Expression Omnibus (GEO) repository under accession number GSE__________. 4C data from the duplicated alleles were reanalyzed from a previous study [51]. The larger size of the ‘Dup’ boxplot shown in Fig. 3f represents the greater amount of fragments used for the quantification of 4C signals.

In order to detect tripartite interactions, one of the 4C libraries was re-sequenced 250bp single-end. The reads were de-multiplexed using fastx_barcode_splitter (http://hannonlab.cshl.edu/fastx_toolkit/) and the viewpoint sequence was removed except the CATG (first cutter sequence) with seqtk (https://github.com/lh3/seqtk). Then they were trimmed for low quality and presence of GATC (second cutter sequence) with cutadapt (cutadapt -q 10 -a GATC)[52]. Next, they were split if a CATG was present. The 5’ part of the split reads (hereafter referred to as mid) and the 3’ part of the split reads (hereafter referred to as third) were mapped independently with bowtie [53](version 0.12.9) on mm10 (bowtie -p 5 -S -k 1 -m 1 -I 0 --best – strata). If the third read did not map, they were split again for CATG and only the first part was now considered as third and mapped. Reads for which the mapping of mid and third were consecutive (undigested situation) were not considered in the analysis. The reads were then pooled according to the mapping position and strand of the mid and the mapping position and strand of the third to remove potential PCR duplicates, resulting in a list of unique tripartite interactions. Each tripartite interaction in the 73.8-74.7 Mb region of the chr2 was assigned to a 20kb bin and the matrix showing the number of different tripartite interactions was plotted.

### CTCF ChIP-seq

A total of 100 mg of distal limbs were dissected from wild type CD1 embryos at E12.5 and fixed during 10 minutes in 1% PFA solution. The fixing reaction was stopped with glycine (0.1M final concentration) and the pellet was washed three times with PBS and stored at -80C. Chromatin extraction and immunoprecipitation were performed using the ChIP-IT High sensitivity kit (Active motif) according to manufacturer specification with slight modifications. Nuclei were extracted and sonicated in 600 ml of Sonication Buffer (50 mM Tris pH=8.0, 1% SDS, 10 mM EDTA) using a Vibra Cell tip sonicator to obtain 200-300 bp average size fragments. Subsequently, 25 mg of sonicated chromatin were diluted ten times in ChIP dilution buffer (20 mM HEPES, 150 mM NaCl, 0.1% NP40) and incubated overnight at 4C with 4 mg of anti-CTCF antibody (Active motif) on a rotating platform. The next day chromatin-antibody complexes were incubated with protein A/G agarose beads during 3 hours at 4C and successively washed following manufacturer instructions. Finally they were eluted and purified by phenol:chlorophorm extraction and precipitation. A total of 20 ng of immunoprecipitated DNA were sequenced with a 50 bp single-end Illumina HiSeq flow cell. Sequenced read were mapped against the mouse GRC38/mm10 genome assembly using the BBCF HTSstation (http://htsstation.epfl.ch and [50]) platform. The dataset were deposited in GEO (accession number GSE__________).

### RT-qPCR

RNA was extracted from pools of micro-dissected limbs or parts thereof, with the Qiagen Tissue Lyser and Qiagen RNeasy Plus kit. 500 ng of RNA was reversed transcribed using random hexamers and Superscript III (Invitrogen). Relative and absolute qPCR were performed with 1 ng of template in technical triplicate. Primers and protocols were described in [54].

## DECLARATIONS

### Acknowledgements

We thank E. Joye for her technical assistance and O. Burri and R. Guiet from the EPFL BioImaging plateform for help with image analysis. We thank S. Boyle and W. Bickmore (MRC, Edinburgh) for designing some of the FISH probes and B. Mascrez, S. Gitto and T.-H. Nguyen Huynh for help with mutant stocks. We thank M. Docquier from the Geneva Genomics Platform, as well as the Bioinformatics Core Facility of the Ecole Polytechnique Fédérale in Lausanne for their assistance in generating and analyzing 4C-Seq and RNA-Seq data. Computations were performed at the Vital-IT (http://www.vital-it.ch) center for high-performance computing of the Swiss Institute of Bioinformatics. We also thank E. Rodríguez-Carballo for comments on the manuscript, as well as F. Darbellay and other members of the Duboule laboratories for discussions and sharing reagents.

## Authors’ contributions

PF: Conceived and carried out the experiments, analyzed the data, wrote the paper.

ML and LD: Analyzed the 4C-seq data, edited the paper

BM: Carried out the experiments, analyzed the data

DN: Provided original data, edited the paper

LB: Provided original data, edited the paper

DD: Conceived the experiments, analyzed the data, wrote the paper.

